# The neuroplasticity of division of labor: worker polymorphism, compound eye structure and brain organization in the leafcutter ant *Atta cephalotes*

**DOI:** 10.1101/2020.03.04.975110

**Authors:** Sara Arganda, Andrew P. Hoadley, Evan S. Razdan, Isabella B. Muratore, James F. A. Traniello

## Abstract

Our understanding of how the design of peripheral sensory structures is coupled with neural processing capacity to adaptively support division of labor is limited. Workers of the remarkably polymorphic fungus-growing ant *Atta cephalotes* are behaviorally specialized by size: the smallest workers (minims) tend fungi in dark subterranean chambers while larger workers perform tasks mainly outside the nest. These strong differences in worksite light conditions are predicted to influence sensory and processing requirements for vision. We found that eye structure and visual neuropils have been be selected to maximize task performance according to light availability. Minim eyes had few ommatidia, large interommatidial angles and eye parameter values, suggesting selection for visual sensitivity over acuity. Large workers had larger eyes with disproportionally more and larger ommatidia, and smaller interommatidial angles and eye parameter values, reflecting peripheral sensory adaptation to ambient rainforest light. Additionally, optic lobe and mushroom body collar volumes were disproportionately small in minims, and within the optic lobe, lamina and lobula relative volumes increased with worker size whereas the medulla decreased. Visual system phenotypes thus correspond to task specializations in dark or light environments and reflect a functional neuroplasticity underpinning division of labor in this socially complex agricultural ant.

## Introduction

Morphology, behavior, and nervous system structure appear to be integrated (Corral-López et al. 2017; Gordon et al. 2017; Iglesias et al. 2018). For example, body size correlates with optical sensitivity and resolution in insect vision (Spaethe and Chittka 2003; Rutowski et al. 2009; Palavalli-Nettimi and Narendra 2018; Taylor et al. 2019). Ants are an ideal model system to examine relationships among behavior, body size, and neuroanatomy because workers have evolved as task specialists in several clades (Hölldobler and Wilson 1990). Scaling patterns of brain size and brain compartment substructure among polymorphic workers, moreover, appear to correspond to foraging ecology and the sensory and cognitive demands of task performance (Gronenberg 2008; Muscedere and Traniello 2012; Gordon et al. 2017). Although olfactory inputs are principal information sources in ants (Hölldobler and Wilson 1990; Czaczkes et al. 2015), vision can be significant in foraging ecology and navigation (Knaden and Graham 2016; Narendra et al. 2017). To home, foragers may use a celestial compass (Wehner 2003; Muller and Wehner 2006), optic flow (Ronacher and Wehner 1995), visual cues and landmark panoramas (Graham and Cheng 2009; Müller and Wehner 2010; Schwarz et al. 2011; Huber and Knaden 2015; Freas et al. 2018), polarized light (Zeil et al. 2014), and canopy patterns (Hölldobler 1980; Beugnon et al. 2005; Rodrigues and Oliveira 2014). Addtionally, visual navigation has been associated with peripheral receptor structure, and primary and higher-order processing brain centers (Gronenberg and Hölldobler 1999; Wehner 2003; Ehmer and Gronenberg 2004; Muller and Wehner 2006; Knaden and Graham 2016), and worker behavioral development may be associated with light-exposure and cued neuroanatomical reorganization in the visual system (Stieb et al. 2010, 2012; Yilmaz et al. 2016).

Ant ommatidia are photoreceptive units that may change in number and structure according to visual needs (Moser et al. 2004; Narendra et al. 2016a). Ommatidia structure affects visual capacity: larger ommatidia enhance light sensitivity, ommatidia number determine image resolution, and lower interommatidial angle improves acuity (Land 1997). Reproductive and worker division of labor in social insects may have selected for differences in compound eye structure (Schwarz et al. 2011; Streinzer et al. 2013). In some ant species, ommatidia number and size scale with worker body size (Menzel and Wehner 1970; Bernstein and Finn 1971; Klotz et al. 1992; Baker and Ma 2006; Schwarz et al. 2011), vary in males and females (Narendra et al. 2016b), and scale differently among polymorphic workers within individual compound eyes and between colonies (Perl and Niven 2016a, b). In bull ants (Greiner et al. 2007; Narendra et al. 2011) and bees (Jander and Jander 2002; Greiner et al. 2004) photoreceptor diameter and eye area increase in nocturnal species in comparison to diurnal species, increasing visual sensitivity. Ommatidia facet diameter is generally smaller in diurnal than nocturnal ants (Narendra et al. 2017), but eye size patterns vary (e.g.(Menzi 1987)).

Visual input from the compound eyes travels to the optic lobes (OL) for primary processing (Gronenberg and Hölldobler 1999). OL investment reflects visual ecology in social insects: in subterranean species, workers are eyeless and OLs are absent whereas diurnal solitary foragers have enormous eyes and their OLs occupy 33% of their brains (Gronenberg and Hölldobler 1999). In paper wasps, queens remain inside the nest and have smaller OLs than foraging workers (O’Donnell et al. 2014), and in the weaver ant *Oecophylla smaragdina*, minor workers nurse brood, rarely leave the nest, and have disproportionally smaller OLs than majors, which forage and defend territory (Kamhi et al. 2017).

The OL is comprised of three regions: lamina (contrast detection), medulla (color vision processing and small field motion), and lobula (color vision processing, wide field motion detection, and shape and panorama construction) (Strausfeld 1989; Gronenberg 2008; Dyer et al. 2011). OL interneurons project to the collar of the mushroom body (MB) calyx for higher-order processing (Gronenberg 2001; Farris 2016). In ants, males, queens and worker brains show differential investment in the medulla, lobula and MB collar (Ehmer and Gronenberg 2004), reflecting different visual ecologies. Peripheral sensory structure should correlate with higher-order processing ability in task-specialized workers, but this linkage is not well understood.

To investigate visual phenotypes within the worker caste, we investigated variation in the structure of the compound eyes, OL, and MB collar in morphologically and behaviorally differentiated workers of the fungus-growing ant *Atta cephalotes*. Worker head widths (HW) range from 0.6 to 4.5mm; this striking polymorphism is associated with the frequency (Wilson 1980) and efficiency (Wetterer 1991; van Breda and Stradling 1994) of leaf harvesting, fungal comb maintenance, brood care, hygienic behaviors, and colony defense. The smallest workers (minims, HW<1.2mm) primarily tend brood and the fungal comb in dark underground chambers (Wilson 1980) whereas media workers (HW=1.2-3.0mm) harvest plant material, traveling along foraging trails beneath rainforest canopy, and the largest workers (majors, HW>3.0mm), are responsible for defense (Powell and Clark 2004; Hölldobler and Wilson 2010). Medias use vision during orientation along trails (Vilela et al. 1987; Vick 2005). Size-variable workers thus have different social roles and experience environments strongly differing in ambient light intensity and visual complexity. It is unlikely that a single eye structure and sensory processing ability has evolved in all workers. We hypothesized that *A. cephalotes* visual system organization is associated with the visual ecology of size-related division of labor and has resulted from selection for adaptive plasticity in ommatidia structure, OL organization and MB collar investment. Specifically, we predicted that workers engaging in within-nest or outside-nest activities (in darkness or light, respectively) would vary in compound eye structure and relative investment in the OL and its constituent parts, and in the MB collar to support the requirements of vision associated to task performance.

## Methods

### Laboratory cultures

Queenright *A. cephalotes* colonies were collected in Trinidad (July 2014) and maintained in a Harris environmental chamber (25°C, 50% relative humidity, 12:12h photoperiod). Artificial nests were constructed from multiple plastic boxes (11cm×18cm×13cm each) connected by plastic tubing (ID=2.5cm). Boxes housing fungal combs had dental stone floors with embedded pebbles to provide air circulation for the fungus. Colonies were fed locally collected leaves free of chemicals and organic produce on alternate days, supplemented with rolled oats, apple, and orange mesocarp.

### Worker size variation and tissue sampling

We sampled polymorphic workers from three colonies (Ac09, Ac20, and Ac21). *A. cephalotes* appears to exhibit triphasic allometry, with three worker size classes (subcastes): minims (HW across the eyes <1.2mm), medias (HW 1.2-3.0mm) and major workers (HW>3.0mm). Each worker was anesthetized on ice and brains were dissected in ice-cold HEPES buffered saline. Compound eyes were removed and stored in 70% ethanol for processing. Because dissection is delicate, we were not always able to preserve the brain and eyes of the same individual.

### Compound eye imaging and structural measurements

Ninety-two intact compound eyes were imaged to create 3D stacks (Fig. 1G) to measure ommatidia number (ON), average ommatidial diameter (D), and interommatidial angle (Δϕ). Eyes were stored in 70% ethanol, washed in 100% ethanol (3×10 min) before mounting. We measured one eye per worker. Extraneous cuticle was removed to allow eyes to lie flat and were then mounted in methyl salicylate between coverslips and imaged using a Fluoview 1 confocal microscope (λ=488nm, step size=3.1μm) with a 20x air objective (NA=0.5, CA=2). Cuticle has natural fluorescence. Eye data were recorded blind to subcaste by randomly assigning identification numbers to eyes. To quantify ommatidia number, image stacks were flattened in ImageJ (Abràmoff et al. 2004) and facets were counted using the Cell Counter plugin. Volume renderings were viewed in Amira 6.0 to verify counts.

**Figure 1:**
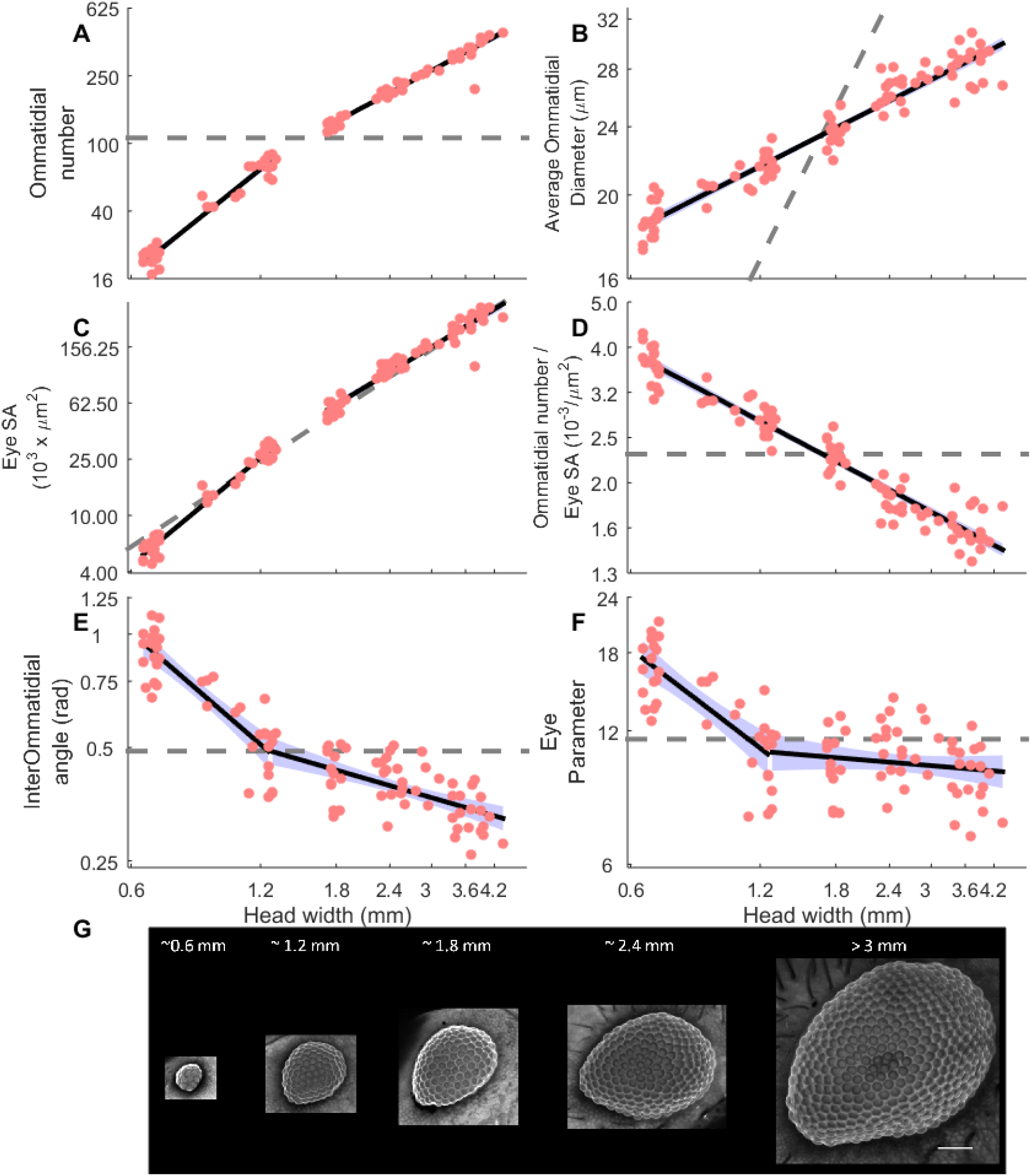
Compound eye structure in polymorphic *A. cephalotes* workers. **A.** Log-log plot of the ommatidia number as a function of worker HW showing a significant change of slope at 1.38mm. **B.** Log-log plot of average ommatidial diameter as a function of worker HW. **C.** Log-log plot of the eye surface area (SA) as a function of worker HW showing a significant change of slope at 1.44mm. **D.** Log-log plot of ommatidia density (ommatidial number/eye SA) as a function of worker HW. **E.** Log-log plot of the interommatidial angle (rad) as a function of worker HW showing a significant change of slope at 1.25mm. **F** Log-log plot of the eye parameter as a function of worker HW (significant change of slope at 1.26mm). **G.** Z-projections of confocal images of eyes from workers with variable HW (scale bar=100μm). A-F: Each pink point represents a single eye. Solid (significantly different from isometry) or dashed (not significantly different from isometry) black lines show linear regression or piecewise linear regressions as appropriate. Purple patches represent 95% confidence intervals of regression lines. Dashed grey lines are the best-fitting isometric regression models.

Mean ommatidial diameter was calculated from the average diameter of 5 or 10 randomly selected ommatidia from each eye. Eye surface area was calculated from the mean ommatidial diameter (surface area=ON×π×[0.5xD]^2^), and ommatidial density (number of ommatidia per surface area unit) was calculated by dividing the number of ommatidia by eye surface area (Yilmaz et al. 2014). To quantify interommatidial angle (Δϕ), image stacks were re-sectioned in the yz plane to obtain a virtual cross section of the eye. ImageJ was used to estimate local eye radius *R* (Schwarz et al. 2011), which with the mean ommatidial diameter (D, in μm) for that eye, interommatidial angle (in radians) could be estimated as Δϕ = *D/R* (Schwarz et al. 2011). Eye parameter (P), which indicates the extent of trade-offs between sensitivity and resolution, was calculated as Δϕ×D (Snyder 1977; Rutowski et al. 2009); lower values of P indicate enhanced acuity, while compromising sensitivity.

### Immunohistochemistry and confocal microscopy

After dissection, brains were placed in 16% Zn-formaldehyde (Ott 2008), fixed overnight at approximately 18°C on a shaker, washed in HBS (6×10 min) and then fixed in Dent’s Fixative (80% methanol, 20% DMSO) for minimally 1h. Brains were next washed in 100% methanol and either stored at −17°C or immediately processed. Brains were washed in 0.1M Tris buffer (pH=7.4) and blocked in PBSTN (5% neutral goat serum, 0.005% sodium azide in 0.2% PBST) at 18°C for 1 hour before incubation for 3 days at room temperature in primary antibody (1:30 SYNORF 1 in PBSTN; monoclonal antibody 3C11obtained from DSHB, University of Iowa, IA, USA). They were washed (6×10 min) in 0.2% PBST and incubated in secondary antibody (1:100 AlexaFluor 488 goat anti-mouse in PBSTN) for 4 days at room temperature. Brains were then washed a final time (6×10 min in 0.2% PBST) and dehydrated in an ethanol series (10min/step, 30/50/70/95/100/100% ethanol in 1x PBS), cleared with methyl salicylate, and mounted on stainless steel slides for imaging.

Sixty-three brains were imaged using an Olympus Fluoview 1 confocal microscope (λ=488nm, step size=3.1μm) with either a 10x air objective (NA=0.3, CA=1), or a 20x air objective (NA=0.5, CA=2). Voxel depth was multiplied by a factor of 1.59 to correct for axial shortening due to mounting in methyl salicylate (Bucher et al. 2000). Brain image stacks were manually segmented using Amira 6.0 and Amira 2019.2 software to quantify neuropil volumes. Given their bilateral symmetry, we segmented one hemisphere per brain, chosen randomly. Our study goal required that only the lamina, medulla and lobula of the OL and MB calyces (separating lip and collar) were segmented separately (Fig. 2A), the rest of the central brain regions and the suboesophageal ganglion were segmented as a whole. Brain data collection was blind to worker HW, although extreme size differences were obvious. Nevertheless, due the randomized coding of brains, subcaste could not be determined with certainty by the annotator. We calculated the volume of the brain hemisphere, the absolute and relative volume of OL (relative to total brain volume), the absolute and relative volume of MB collar (relative to total brain volume), and the relative volumes of OL subregions (relative to total OL volume).

**Figure 2:**
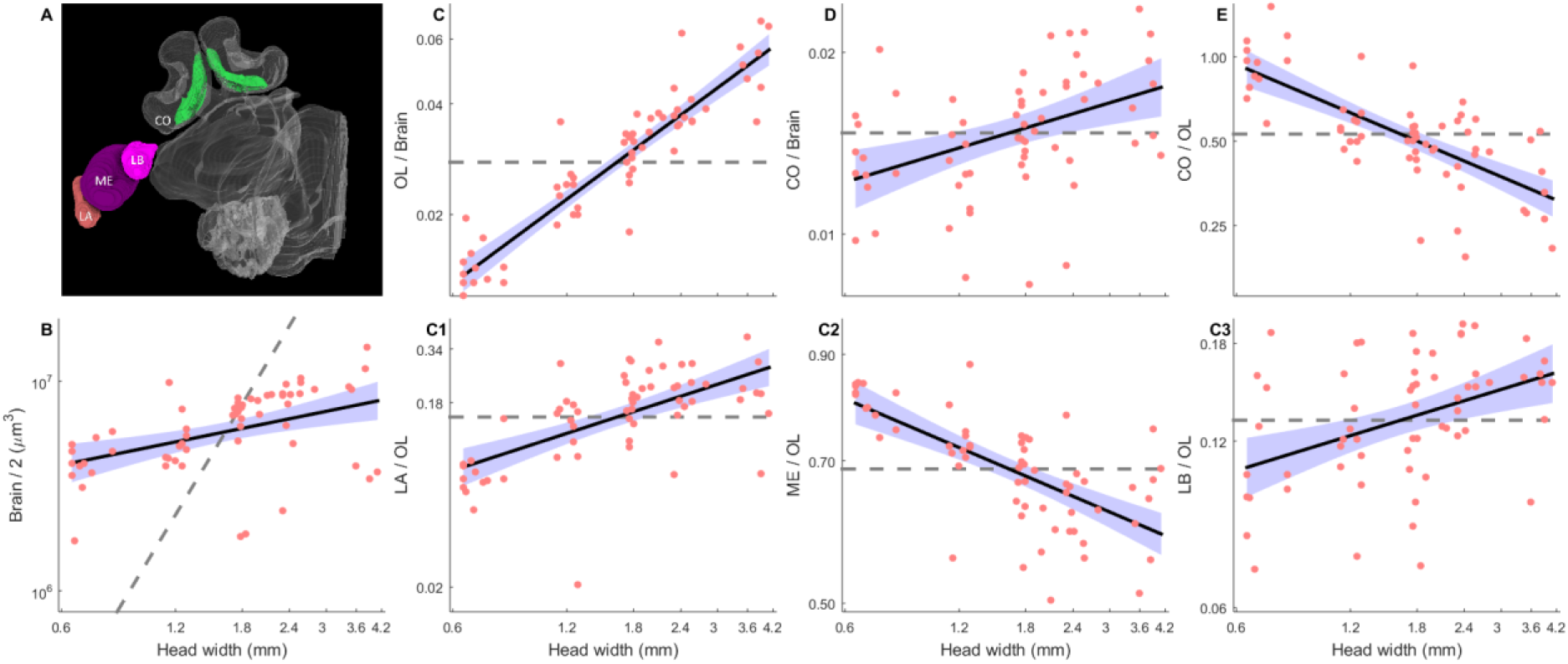
Volumes of polymorphic *A. cephalotes* worker brains and brain compartments. **A.** 3D reconstruction of the brain hemisphere of an *A. cephalotes* worker (HW ~4mm). **B.** Log-log plot of hemisphere brain volume as a function of worker HW. **C.** Log-log plot of relative OL volume as a function of worker HW. **C1.** Log-log plot of relative volume of OL lamina as a function of worker HW. **C2.** Log-log plot of relative volume of OL medulla as a function of worker HW.**C3.** Log-log plot of relative volume of OL lobula as a function of worker HW.**D.** Log-log plot of relative MB collar volume as a function of worker HW. **E.** Log-log plot of MB collar: OL volume ratio as a function of worker HW. B-E: Legend as in figure 1.

### Statistical evaluation

Statistical evaluations were performed in R (version 3.3.0, Team 2016) using the ‘segmented’ package to analyze eye and brain metric scaling (V. R. M. Muggeo 2008). To assess allometries in eye structure and brain volumes in relation to worker size, least-square means regression was used on log10-transformed values to estimate *a* and *b* in the scaling equation *y*=*aM^b^*, as *log*10(*y*)=*log*(*a*)+*b*×*log*(*M*). To test the null hypothesis (H0) of isometry, a separate linear model was calculated and tested against different slope values depending on the metric. The slope for H_0_ were *b*=0.0 (linear vs. constant values), *b*=1.0 (linear vs. linear), *b*=2.0 (linear vs. surface area) and *b*=3.0 (linear vs. volume) (Kaspari and Weiser 1999).

The Davies test was used to determine if there is a statistically significant change in slope or a ‘breakpoint’ in a linear relationship (Davies 2002). We observed that the significance of some changes in slope depended on a single data point; therefore, we accepted the change in slope only if its significance was always below 0.05 when removing any point from the dataset. The ‘segmented’ package was further used to estimate the location of the breakpoint. If the Davies’ test revealed two piecewise linear relationships in a scaling relationship, least-square means regression was calculated and tested against isometry independently.

To further explore whether increased investment in primary visual neuropil might have an impact in higher-order visual processing neuropil, we assessed allometry in the ratio of volumes of the optic lobes and MB collar according to HW. We also calculated a least-square means regression on log10-transformed values and tested against isometry (*b*=0.0).

## Results

### Eye Structure

The eyes of media and major workers had significantly more ommatidia than minims (Fig. 1A) and showed a significant change in the scaling of ommatidia number and worker size (Davies test, p<0.001) at a HW of 1.38mm (95% CI: 1.20 to 1.58mm). Piecewise linear models calculated for both slopes were significant (p<0.001, Multiple R^2^=0.989) with a slope shift from 2.03 (95% CI: 1.91 to 2.13) to 1.35 (95% CI: 1.25 to 1.45). Piecewise linear models were also significantly different from isometry (*b*=0; p<0.001). Media and major workers also had larger ommatidial diameter (Fig. 1B). The relationship between ommatidial diameter and worker size showed no significant breakpoint (Davies test, p>0.5), and these variables were significantly correlated (F_(1,90)_=1217, p<0.001, R^2^=0.93). The slope of the regression line was 0.25 (95% CI: 0.24 to 0.27), which was significantly different from isometry (*b*=1.0; F_(1,90)_= 1217, p<0.001).

Compound eye size (total eye surface area) was correlated with HW (Fig. 1C). Davies’ test showed a significant change in the scaling of total eye size and worker size (p<0.001) at a HW of 1.44mm (95% CI: 1.20 to 1.72mm). Piecewise linear models calculated for both slopes were found to be significant (p<0.001, Multiple R^2^=0.988) with a slope shift from 2.52 (95% CI: 2.36 to 2.68) to 1.84 (95% CI: 1.70 to 1.97). Piecewise linear models were significantly different from isometry (*b*=2.0; p<0.001), although the effect size was small.

The density of ommatidia decreased with HW (Fig. 1D). The relationship between the density of ommatidia and worker size showed no significant breakpoint (Davies test, p>0.5 and these variables showed a significant correlation (F_(1,90)_=1217, p<0.001, R^2^=0.93). The slope was −0.50 (95% CI: −0.53 to −0.48), also significantly different from isometry (*b*=0.0; F_(1,90)_= 1217, p<0.001).

Interommatidial angle decreased as worker size increased (Fig. 1E). Davies’ test for a change in slope showed a significant change in the scaling relationship between interommatidial angle and worker size (p<0.001) at a HW of 1.25mm (95% CI: 1.03 to 1.51mm). Piecewise linear models were calculated for both slopes and found to be significant (p<0.001, Multiple R^2^=0.84), with a slope shift from −0.98 (95% CI: −1.21 to −0.76) to −0.33 (95% CI: −0.43 to − 0.23). Piecewise linear models were also significantly different from isometry (*b*=0; p<0.001).

Eye parameter decreased with worker size in minims (Fig. 1F). Davies’ test showed a significant change in the scaling relationship between eye parameter and worker size (p<0.001) at a HW of 1.26 mm (95% CI: 1.01 to 1.57 mm). Piecewise linear models were calculated for both slopes and found to be significant (p<0.001, Multiple R^2^=0.588) with a slope shift from - 0.73 (95% CI: −0.95 to −0.51) to −0.07 (95% CI: −0.20 to 0.05). The first segment of the piecewise linear models was found significantly different from isometry (*b*=0; p<0.001), but the second segment was not (*b*=0; p=0.205).

### Brain Structure

Larger workers had significantly larger brains (Fig. 2B). The relationship between brain volume and worker size showed no significant breakpoint (Davies test, p>0.05) and these variables showed a significant positive correlation (F_(1,61)_=13.91, p<0.001, R^2^=0.18) with a slope of 0.37 (95% CI: 0.17 to 0.56) significantly different from isometry (*b*=3.0; F_(1,61)_=13.91, p<0.001). We found greater OL investment in media and major workers (Fig. 2C). Relative OL volume and worker size showed a significant positive correlation (F_(1,61)_=271.2, p<0.001, R^2^=0.81) with no significant breakpoint (Davies test, p=1). The slope of the regression line was 0.76 (95% CI: 0.67 to 0.85) and significantly different from isometry (*b*=0.0; F_(1,61)_=271.2, p<0.001).

Within the OL, relative investment in lamina increased with worker size (Fig. 2C1). Relative lamina volume and HW showed a significant positive correlation (F_(1,61)_=37.77, p<0.001, R^2^=0.37) with no significant breakpoint (Davies test, p>0.05). The slope of the regression line was 0.64 (95% CI: 0.43 to 0.85) and significantly different from isometry (*b*=0.0; F_(1,61)_=37.77, p<0.001). Relative investment in the medulla, in contrast, decreased with worker size (Fig. 2C2). Relative medulla volume and HW showed a significant negative correlation (F_(1,61)_=51.1, p<0.001, R^2^ of 0.45) with no significant breakpoint (Davies test, p>0.05) and a slope of −0.17 (95% CI: −0.21 to −0.12), significantly different from isometry (*b*=0.0; F_(1,61)_=51.1, p<0.001). Finally, as for the lamina, relative investment in the lobula increased with worker size (Fig. 2C3). Relative lobula volume and HW showed a significant positive correlation (F_(1,61)_=13.43, p<0.001, R^2^=0.17) and no significant breakpoint (Davies test, p=0.362). The slope of the regression line was 0.21 (95% CI: 0.10 to 0.32) and significantly different from isometry (*b*=0.0; F_(1,61)_=13.43, p<0.001).

Media and major workers also invested relatively more in the MB collar (Fig. 2D). Relative collar volume and worker size showed a significant positive correlation (F_(1,61)_=13.03, p<0.001, R^2^=0.16) and no significant breakpoint (Davies test, p=0.42). The slope of the regression line was 0.19 (95% CI: 0.08 to 0.29), which was also significantly different from isometry (*b*=0.0; F_(1,61)_=13.03, p<0.001). Despite investing more in the MB collar, larger workers had a lower collar:OL volume ratio (Fig. 2E). The relationship between this ratio and worker size showed a significant negative correlation (F_(1,61)_=67.17, p<0.001, R^2^=0.52), no significant breakpoint (Davies test, p=0.69), and a slope of −0.57 (95% CI: −0.71 to −0.43), significantly different from isometry (*b*=0.0; F_(1,61)_=67.17, p<0.001).

## Discussion

Social insect compound eyes and visual information processing neuropils enable adaptive behavioral performance according to cognitive challenges of navigation and ambient light levels (Jander and Jander 2002; Mares et al. 2005; Kapustjanskij et al. 2007; Warrant 2008; Narendra et al. 2011, 2016a; Streinzer et al. 2013; Yilmaz et al. 2014; Bulova et al. 2016). Eye size and ommatidia number correlate with worker size in ants, including polymorphic species, and may be associated with task performance (Menzel and Wehner 1970; Klotz and Reid 1992; Schwarz et al. 2011). In polymorphic *A. cephalotes* workers, in which task performance is strongly correlated with body size-related division of labor, differences in worksite sensory ecology appear to select for visual system polyphenism.

### Division of labor and eye structure in A. cephalotes

*A. cephalotes* workers perform tasks in the complete darkness of fungal comb chambers and in the filtered light epigaeic environment beneath rainforest canopy. Media workers forage day and night (Cherrett 1968), and use trail pheromones as well. Although olfaction appears to be the dominant sensory modality for foraging in many ants, visual information facilitates trail-following in *Atta laevigata* [69], and other ant species (Beugnon and Fourcassié 1988) alter their use of chemicals or vision depending on light conditions. In *A. cephalotes*, improved forager visual ability may enable flexibility in the use of orientation cues and social signals as ambient light levels change.

Minims tend fungi deep underground, medias harvest leaves from their habitat and labor inside the nest, and majors appear to exclusively perform defense and trail maintenance outside. We hypothesized that eye structure variation among subcastes would reflect adaptation to worksite light availability and visual demands for task performance. We expected the eyes of minims to structurally enable light sensitivity over resolution, whereas larger worker eyes were predicted to favor spatial resolution over sensitivity. It is unclear how minims make use of visual information and what level of spatial resolution and sensitivity is needed to work effectively on the fungal comb. Minims, however, also perform some tasks outside the nest, “hitchhiking” on transported leaves during day and night to defend against fly parasites (Linksvayer et al. 2016). We found that the number and size of ommatidia and eye surface area were significantly smaller in minims (Fig. 1A-C), suggesting less capacity to capture light and less reliance on vision to perform their tasks. The larger ommatidia of media and major workers (Fig. 1B) indicate greater sensitive to light. However, ommatidia size increased hypometrically with body size: although the ommatidia of minims were the smallest, relative ommatidia size was greater in minims than in medias and majors. This may enable minim worker eyes to collect more light than expected from their size, suggesting adaptation to darkness and light (Greiner 2006; Yilmaz et al. 2014). Alternatively, the small size of minim worker eyes and ommatidia may be due to a body size constraint: eye size is as large as developmentally possible to ensure at least a marginal ability to capture light, which may be needed for parasitic fly defenses. Minims also showed a higher density of ommatidia than media and majors (Fig. 1D). Interommatidial angle decreased with worker size, indicating higher visual acuity in larger workers (Fig. 1E). Eye parameter values (Fig. 1F) were significantly higher for minims, but for larger workers, values were lower and not correlated with size. This suggests minim worker eyes are adapted to enhance sensitivity rather than acuity, whereas larger worker eyes structure have been selected for sensitivity and acuity. Higher acuity is adaptive outside the nest, as it allows resolving more distal objects. The significant breakpoints (HW 1.0-1.8mm) found in the linear regressions for ommatidia number, eye surface area, interommatidial angle and eye parameter (Fig. 1A,C,E,F) suggest structural changes to accommodate the body size-associated transition between inside and outside nest division of labor in *A. cephalotes*. Comparisons of eye structure between diurnal, cathemeral, and nocturnal ant (Greiner et al. 2007; Narendra et al. 2013; Yilmaz et al. 2014; Ogawa et al. 2019) and bee species (Greiner et al. 2004) are generally consistent with our predictions.

Although eye structure determines light sensitivity and visual acuity, other anatomical, physiological, and behavioral adaptations modify visual abilities: variations in the size of rhabdomers (Greiner et al. 2004; Gonzalez-Bellido et al. 2011; Narendra et al. 2017), microsaccadic rhabdomere contractions and microvilli refractory time (Juusola et al. 2016), or pupillary systems mediated by pigment ommatidial cells (Narendra et al. 2013, 2016a) Such visual adaptations in *A. cephalotes* polymorphic workers remain to be studied.

### Division of labor and visual neuropil size and structure

In ant species characterized by morphological differentiated subcastes, workers are predicted to vary neurobiologically to support the sensory demands of specialized tasks (Muscedere and Traniello 2012; Kamhi et al. 2015; Gordon et al. 2019). If metabolic costs associated with the production and maintenance of brains is high, then selection should favor the reduction of neuropil size (e.g.(Aiello and Wheeler 1995; Niven and Laughlin 2008)). We found that brain volume increased with worker size, but larger workers had brains smaller than expected from their body size (Fig. 2A). We expected *A. cephalotes* workers would invest differentially in brain compartments due to their body size-related task repertoires, and found that larger workers had larger eyes and an allometric increase in OL volume (Fig. 2B). Conversely, some diurnal moths have smaller eyes but larger optic lobes than nocturnal species (Stöckl et al. 2016), a pattern also found in analogous brain region in teleost fishes (Iglesias et al. 2018). *A. cephalotes* OL were disproportionally larger in larger workers, and consistent with our prediction, minims showed disproportionately less OL investment. This suggests a task-related increasing need for primary visual information processing in larger workers.

Our analysis revealed that lamina, medulla and lobula increased with worker size (Fig. S1), maybe due to higher exposure to light in larger workers active outside the nest (as in (Yilmaz et al. 2016)). Within the OLs, larger workers possessed disproportionally larger lamina and lobula, but a disproportionally smaller medulla (Fig. 2C1-C3). These OL subregion allometries suggest that minims might be better at detecting small-field motion whereas larger workers might be better at processing contrast, wide-field motion, shape, and panorama information. This neuroplasticity seems to adaptively support *A. cephalotes* task specialization inside and outside the nest. We also found a disproportional investment in the MB collar in larger workers (Fig. 2D). Enlarged MBs in social hymenopterans might be the result of ancestral neuroanatomical adaptations to process novel visual information [60]. This evolutionary scenario across phylogenetically diverse ant species appears to be reflected in *A. cephalotes* subcastes that vary in visual ecologies. Our results suggest that the increased need for visual cognition in larger workers is greater for primary processing than for higher-order processing. In *Myrmecia* species, nocturnal workers invested relatively less in OL but relatively more in the MB, including the collar, than diurnal workers (Sheehan et al. 2019). Our results showed that minims had the highest collar:optic lobe ratio (Fig. 2E), apparently as an adaptation to performing tasks in darkness. Collaterally, studies of gene expression differences in whole brains of *A. cephalotes* subcastes revealed a significant worker size-related increase in the level of a gene associated with rod cell development, mirroring the higher demand for visual acuity and larger eye structures in larger workers (Muratore et al., unpublished data). This trend was also true for a gene associated with growth factor activity, potentially contributing to the allometric OL enlargement and other brain regions.

## Conclusions

We found optical and neural plasticity are associated with the complex agrarian division of labor of *A. cephalotes* workers. Previous studies describe differences in eye structure (Menzel and Wehner 1970; Bernstein and Finn 1971; Klotz et al. 1992; Baker and Ma 2006; Schwarz et al. 2011) or visual neuropil investment (O’Donnell et al. 2018). Our results advance our understanding of ant visual system functionality by demonstrating caste-related compound eye and brain plasticity that has evolved in response to worksite light levels. Worker polymorphism has been shown to be correlated with patriline in the several leafcutting ant species (Hughes et al. 2003; Evison and Hughes 2011), suggesting a potential link between genetic variation and the neuroanatomical patterns described here. Division of labor underpinning the fungicultural habits of *A. cephalotes* appears to have played an important selective role in worker visual system evolution. Worker behavior in this species, however, depends on visual and olfactory information that likely varies with the cognitive requirements of tasks. The influence of these factors on the spatial resolving power and sensitivity of eyes and macroscopic and cellular structure of *A. cephalotes* brains requires further study.

## Author’s contributions

APH and ESR prepared brains for imaging and registered images; APH, ESR and SA analyzed neuropil volumes; ESR initiated and designed the eye metric imaging protocol, registered eye images, and measured eye structure; APH and SA performed statistical analysis; APH collected ant colonies; IBM analyzed confocal images and contributed to manuscript content; ESR and APH prepared the first draft of the manuscript; JFAT, APH, ESR, and SA conceptualized and designed the study. JFAT secured funding for the study; JFAT and SA wrote the manuscript. All authors edited and approved the final content of the manuscript.

## Acknowledgments

We are very grateful for the assistance of Dr. Darcy Gordon (study design, scaling analysis, and statistics), Dr. Christopher Starr and Ricardo “Monkey” Pillai (colony collection), Dr. Todd Blute (confocal training), Dr. Alfonso Pérez-Escudero (data analysis and interpretation), and Dr. J. Frances Kamhi (insightful comments on the manuscript). This research was supported by National Science Foundation grant IOS 1354291 to JFAT, a Marie Skłodowska-Curie Individual Fellowship BrainiAnts-660976 and Ayudas destinadas a la atracción de talento investigador a la Comunidad de Madrid en centros de I+D to SAC, the Undergraduate Research Opportunities Program at Boston University to ESR, and a National Science Foundation Graduate Research Fellowship (DGE-1247312) to APH.

