## Supplementary material for "The neuroplasticity of division of labor: worker polymorphism, compound eye structure and brain organization in the leafcutter ant *Atta cephalotes*": Fig. S1

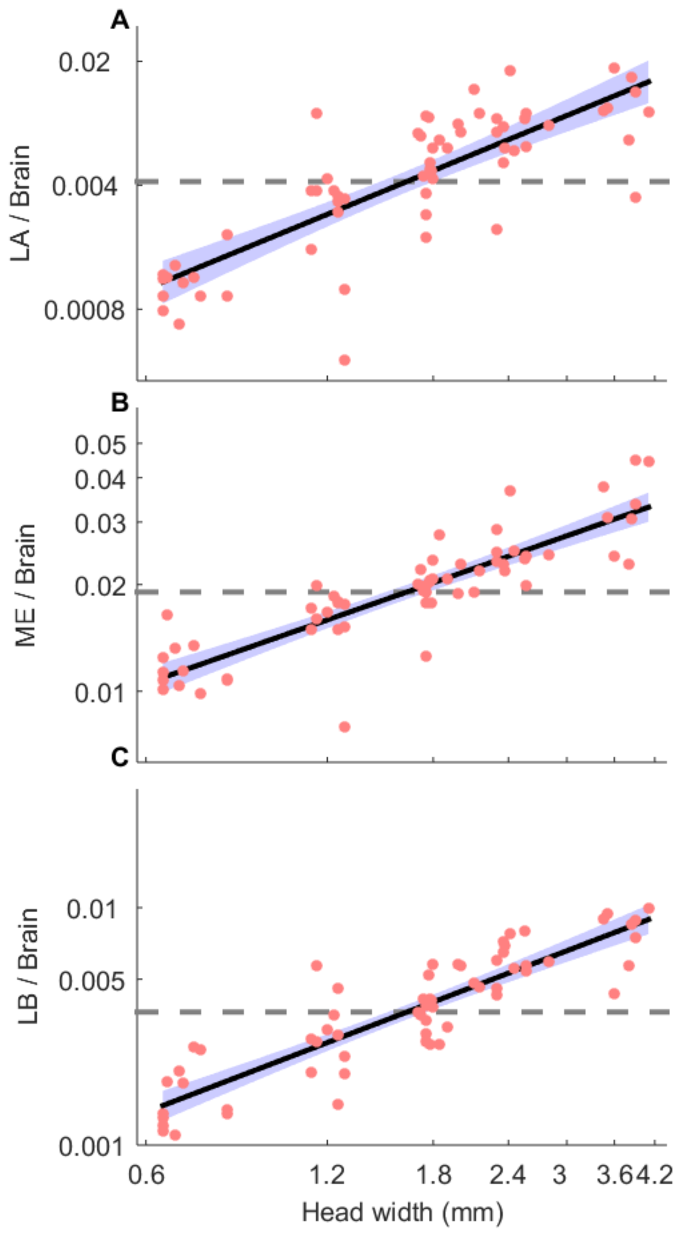


**Figure S1: Brain volumes of polymorphic *A. cephalotes* workers. A.** Log-log representation of lamina volume (LA) relative to hemisphere brain volume (Brain) as a function of worker HW. Slope of 1.40 (95% CI: 1.14 to 1.66) significantly different from isometry ($b$=0.0; $F_{(1,61)}$= 116.3, $p<0.001$). **B.** Log-log representation of medulla volume (ME) relative to brain volume as a function of worker HW. Slope of 0.60 (95% CI: 0.51 to 0.68) significantly different from isometry ($b$=0.0; $F_{(1,61)}$= 180, $p<0.001$). **C.** Log-log representation of lobula volume (LB) relative to brain volume as a function of worker HW. Slope of 0.97 (95% CI: 0.84 to 1.10) is significantly different from isometry (*b*=0.0; $F_{(1,61)}$= 213.3, $p<0.001$). Each pink point represents a single eye. Solid black lines show linear regression or piecewise linear regressions as appropriate. Purple patches represent 95% confidence intervals of the regression lines. Dashed grey lines are the best-fitting isometric regression models.
